# Sunfish as zooplankton control agents to improve yields of wastewater-cultivated algae

**DOI:** 10.1101/2023.12.22.573129

**Authors:** Chase J. Rakowski, Mathew A. Leibold, Schonna R. Manning

## Abstract

Wastewater algal cultivation (WAC) in outdoor raceway ponds is a promising means of removing polluting nutrients from water while producing biomass feedstock for products like fertilizer or biofuel. However, algal cultivation in open ponds is often plagued by underproduction due to contamination by zooplankton. The use of biological controls, i.e., natural predators to control zooplankton, was evaluated as a potential low-cost solution in this study. Eighteen pilot-scale raceway ponds were inoculated with cultured algae and naturally-occurring zooplankton; bluegill sunfish (*Lepomis macrochirus*) were tested as biological control agents to suppress zooplankton. We installed cages in the ponds to protect the fish from the paddlewheels and control the area the fish could access. The cage either surrounded the paddlewheel – in which case the fish had access to the remaining area – or the cage was opposite the paddlewheel, in which case the fish had access to the smaller area inside the cage. We also implemented fishless controls. After 19 days, ponds with cages surrounding the paddlewheel had 93% more zooplankton, but the fish had reduced zooplankton by 89% on average across cage treatments. Cage placement did not affect algal biomass, but ponds with fish had 45% higher algal dry weight and 84% higher algal biovolume. However, no differences in nutrient levels were recorded among treatments after 19 days. We concluded hardy zooplanktivorous fish represent a promising means of boosting algal production in WAC, although further research is needed to determine whether the change in algal biomass could eventually translate to improved nutrient removal.

## Introduction

Microalgal cultivation is a promising means of producing environmentally-friendly alternative fertilizers, animal feeds, fuels, and various higher-value products (Benemann 2013). Some reasons for this potential are that microalgae are extremely productive, they produce useful compounds such as omega-3 fatty acids, and they do not compete with food crops for arable land (Chisti 2007). Besides being cultured for natural products, microalgae can also be used to clean wastewater since they take up the excess nutrients that would otherwise act as pollutants and convert them into biomass. Wastewater here can either refer to partially treated municipal wastewater or hypereutrophic runoff from fertilized fields (e.g. industrial agricultural fields or golf courses). Wastewater algal cultivation (WAC) systems represent an especially promising approach of converting pollution into useful products, eliminating the need for costly fertilizer to grow the algae (Christenson and Sims 2011). The most economical way to do this seems to be to allow wastewater to flow into outdoor circulating ponds termed “raceway ponds” where microalgae are allowed to grow, taking up the excess nutrients as they multiply. Then a fraction of the water, along with suspended biota, is periodically transferred to a settling unit where the algae settle to the substrate for harvest (Park and Craggs 2010). However, yields of cultivated microalgae including in wastewater systems are generally far lower than what is theoretically possible, creating an obstacle to the adoption and operation of WAC (Benemann 2013). One reason is that as open systems, outdoor raceway ponds are inevitably colonized by various aquatic organisms, leading to the assembly of a plankton community including herbivores that reduce algal yields (Smith et al. 2010).

Many herbivorous zooplankton lay resting eggs which disperse via wind, rain, and flying animals (Havel and Shurin 2004), so they often colonize algae ponds and trigger large reductions in algal biomass (Smith et al. 2010; Park et al. 2011). Mechanical and chemical methods of pest control can be used to control nuisance zooplankton but represent a significant recurring cost, may cause collateral damage to the aquaculture system and downstream environment, and are often ineffective at controlling the resistant zooplankton resting eggs (Montemezzani et al. 2015). Rather than using a pesticide that would have to be reapplied indefinitely, or simply draining and re-inoculating the pond – neither option is economical – biological control offers a way to establish long-term self-sustaining pest management (Montemezzani et al. 2015). Biological control methods have the potential to be effective, low cost, and low risk, as long as the biocontrol agents are self-sustaining and native (Smith and Crews 2014). However, scarce research has been done on biological control as a means of improving yields of microalgae. While a few review papers have discussed the potential for biological pest control in algae ponds, and listed taxa that may have the potential to function as biological control agents (Smith et al. 2010; Smith and Crews 2014; Montemezzani et al. 2015), we are only aware of one published experiment that tested a biological control method for improving algal yields (Sturm et al. 2012). This paper found that mosquitofish enhanced algal yields in still wastewater settling ponds, leaving open the question of biological control efficacy in raceway ponds whose paddlewheels could pose a danger for larger organisms.

Here we test whether bluegill sunfish (*Lepomis macrochirus*) can improve the production efficiency of raceway WAC systems – which is to say increasing algal production and uptake of polluting nutrients – by suppressing zooplankton. By installing cages in miniature raceway ponds, we protected the sunfish from the rotating paddlewheels and controlled the area they could access. We predicted that the fish would reduce zooplankton biomass. Additionally, when outside the cage and given access to more area and the tank bottom, we predicted that the fish would also reduce more mobile and benthic invertebrates, at the expense of zooplankton suppression. Managers may wish for the fish to primarily consume zooplankton rather than benthic or more mobile invertebrates, which likely do not pose a threat to phytoplankton production and, in the case of predaceous insects, could help suppress zooplankton. Furthermore, we predicted that according to trophic cascade theory, each successive lower trophic level would be affected in the opposite way from the trophic level above (Carpenter et al. 1985). Therefore, we predicted algal biomass would be lowest without fish, intermediate with fish outside the cage, and highest with caged fish, and that dissolved nutrient concentrations would be affected in the same qualitative way as zooplankton biomass: highest without fish, intermediate with fish outside the cage, and lowest with caged fish.

## Methods

### Raceway ponds

In an unshaded field in Brackenridge Field Laboratory, we constructed miniature raceway ponds using 120 (l) x 68 (w) x 32 (d) cm round-end 230 L plastic stock tanks lined with white shower curtain liners that were weighted down with three white tiles. We placed wooden supports underneath to ensure all tanks were level. In the center of each tank we placed a clear acrylic baffle to allow circular (“raceway”) flow. A clear acrylic paddlewheel supported by a white plastic rod connecting three or four ponds continuously circulated the water at 6±1 cm/s. To provide the fish with protection from predators, we covered 29% of each tank with black mesh with 2 cm^2^ holes on the side opposite the paddlewheel. On May 5, as air temperatures increased after the experiment had begun, we replaced the mesh with an equivalent area of black weed blocker to provide shade and reduce water temperatures (Fig. 1).

**Figure 1.**
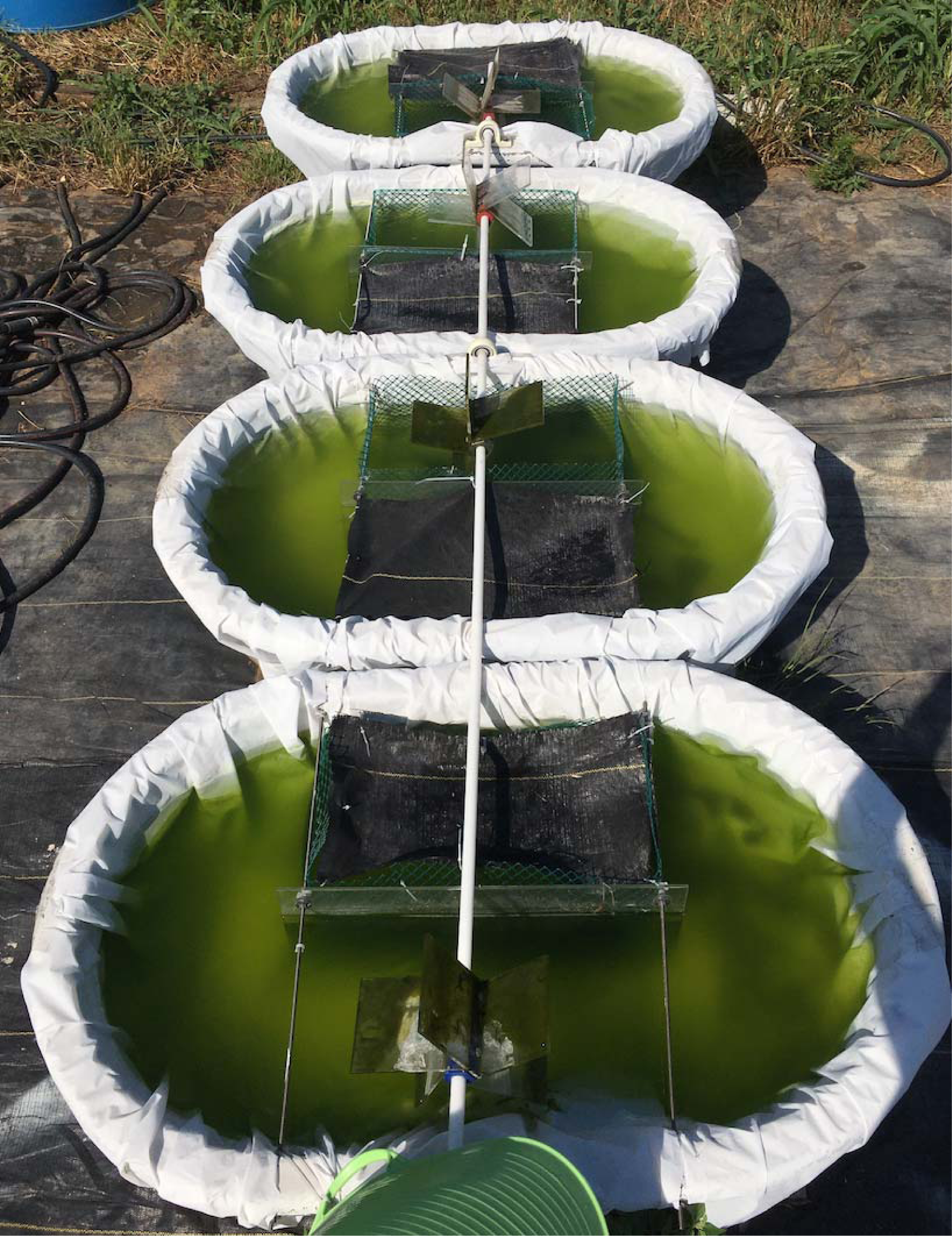
A row of miniature raceway ponds. The pond in the foreground (bottom of photo) has a cage opposite the paddlewheel, whereas the others have a cage surrounding the paddlewheel.

On February 21, 2022 we filled the tanks with 230 L Austin municipal water which is rich in minerals including 15 mg/L Mg^2+^ and 12 mg/L Ca^2+^. We added 132 µL/L Amquel (hydroxymethanesulfonate) to eliminate chloramine and ammonia. Then to create synthetic wastewater, we added 370.761 mg/L (NH_4_)_2_SO_4_, 225.2 mg/L urea (CH_4_N_2_O), 134.79 mg/L KH_2_PO_4_, and 85.45 mg/L K_2_HPO_4_ in dissolved form. We also added 10 µL/L Advanced Nutrients Jungle Juice Micro, which contains 5% Ca, 4.7% NO_3_-N, 0.3% NH_4_-N, 1% K_2_O, 0.1% chelated Fe, 0.05% chelated Mn, 0.015% chelated Zn, 0.01% B, 0.01% chelated Cu, 0.0008% Mo, and 0.0005% Co derived from calcium nitrate, ammonium nitrate, potassium nitrate, potassium borate, iron DTPA, iron EDDHA, iron EDTA, manganese EDTA, zinc EDTA, copper EDTA, ammonium molybdate, and cobalt nitrate. The resulting media had a pH of 7.16, N:P ratio of 10.0, 131.5 mg/L total Kjeldahl N, and 1 mg/L NO_3_-N. The composition marked by high ammonium, organic nitrogen, and phosphates but moderate nitrates was intended to be broadly representative of secondary municipal and agricultural wastewater (US EPA 2011). Leading up to the experiment we replaced evaporated media with fresh media. Then during the experiment we performed 12.5% media exchanges every 2 days to imitate continuous cultivation with a 16-day holding time.

### Organism collection

After filling the tanks with media, we added 13 mL dense steady state culture of UTEX 1230 *Chlorella sorokiniana* to each tank. We then collected mixtures of phytoplankton and zooplankton from other small water bodies in Brackenridge Field Laboratory, mixed them, and added equal volumes to each tank. We similarly added equal volumes of mixtures of zooplankton and *Neocorixa snowi* (water boatmen), as well as 15 *Streptocephalus* (fairy shrimp), collected from the A. E. Wood Texas Parks and Wildlife Fish Hatchery in San Marcos, TX to each tank on April 25. Succession of the pond communities was allowed to proceed until the start of the experiment.

We collected juvenile bluegill sunfish (*L. macrochirus*) from a breeding population in a small manmade pond continuously fed by municipal water near the experimental site in Brackenridge Field Laboratory by deploying a live fish trap for <18 hours on April 20-21 and 24-25. We returned adults and juveniles >9 cm standard length to the pond, then transported the remaining juveniles (6-9 cm standard length) to the experiment site in buckets filled with pond water (<15 minutes). Upon arrival we acclimated the fish for >1 day in 300 L tanks filled with a 1:1 mixture of the artificial wastewater media and municipal water with Amquel added, roughly representing a halfway point between the water chemistry of their native pond and the experimental raceway ponds.

### Experimental design & sampling

We implemented three treatments of the area fish could access in tanks, which we expected to correlate with the degree of predation especially on more mobile invertebrates. In each tank between the baffle and one wall we placed a 0.2 m^2^ (29% of total pond area) cage made of plastic poultry mesh with 3 cm^2^ holes, eight on the side opposite the paddlewheel and ten surrounding the paddlewheel. This hole size allowed invertebrates but not fish to pass through. Due to the curvature of the tank bottom, only one cage edge was flush with the bottom, making it more difficult for caged fish to access the benthos. To six of the cages opposite the paddlewheel, we added one juvenile sunfish each on April 29. At the same time we also added a juvenile sunfish to six of the tanks with cages surrounding the paddlewheel, outside the cage so that the fish had access to the remaining 0.48 m^2^ of the tank area. The remaining tanks, two with cages opposite the paddlewheel and four with cages surrounding the paddlewheel, served as fishless controls. Thus, more precisely there were two fully crossed treatments: fish presence and cage placement, allowing us to separate any effect of the proximity of the cage to the paddlewheel from the effect of fish access. The treatments were assigned to tanks randomly in space.

We visually assessed prey densities in the removed media and when densities fell below visually detectable levels, we supplemented all tanks equally with zooplankton (May 12) or *Streptocephalus* (April 30, May 8). We measured water temperature and pH with a sonde daily. On days when the high (air) temperature was predicted to exceed 32 °C, we added enough ice to cool the water to 30 °C between 12 and 2 PM. When the high (air) temperature was predicted to reach or exceed 38 °C, we monitored the water temperature with a sonde following the initial ice addition and if the temperature was approaching 33 °C, we added cool water if needed to keep the temperature below 33 °C. We visually assessed the health of each fish daily and replaced any moribund or deceased fish with a fish from the acclimation tank. Moribund fish were humanely euthanized following IACUC animal use protocol # AUP-2021-00240, and all other methods were conducted according to this AUP.

On March 11, we collected live algal samples for isolation and identification via DNA barcoding. Then on May 17, we collected samples for preservation and chemical analyses. We collected 1 mL media from each tank and preserved it in 2.5% formalin for microscopy analysis of algae. To sample the invertebrate community, we used a 6-L bucket to collect 12.5% of the media in each raceway pond (28.75 L) from the whole water column including the benthos. Then we filtered this sample with 45 µm mesh and preserved the retained material in 70% ethanol. We kept 1 L of the filtered liquid from each pond for chemical analyses and maintained it on ice for <10 hours. We then centrifuged it to separate the algae from the media, and dried the algae in a drying oven at 80 °C for 12 hours and placed it in a desiccator for 30 days. We added 2 µL/mL concentrated (18 M) sulfuric acid to the media samples to bring the pH under 2.0 and stored them in a refrigerator at 4 °C for 28 days.

### Analysis of algae and invertebrates

For the common algal morphospecies which could not be identified visually, we isolated individual cells from the March 11 live samples with a capillary pipette and cultured them in Modified Bold’s 3N medium for >3 weeks. Then we extracted the DNA using a GENECLEAN® kit and ran PCR using both forward and reverse 23S and ITS2 primers. We checked all PCR products using gel electrophoresis and cleaned them using a ThermoFisher ExoSAP-IT™ kit. Finally, we sent the PCR products to Eton Bioscience for Sanger sequencing. Sequence reads were visualized and edited, e,g. deleting N’s, using FinchTV (FinchTV^®^ 1.5.0, Geospiza, Inc., Seattle, WA), then matched with published sequences on NCBI using blastn.

To estimate biovolumes of phytoplankton taxa, we calculated densities of each morphospecies with a hemocytometer. We counted at least 200 cells of the most common morphospecies or 900 nL – whichever came first – and at least 600 nL for less common taxa. We captured several micrographs of each morphospecies from various tanks and sampling dates, and used ImageJ to measure the cell dimensions of at least 40 cells for the common morphospecies, at least 10 cells for the rarer morphospecies, and as many as could be found for the rarest morphospecies (Schneider et al. 2012). We then calculated the biovolume of each cell using geometric approximations (Online Resource 1 Table S1).

To estimate biomass of invertebrate taxa, we identified, counted, and measured individuals in subsamples such that for each taxon, we counted at least 200 individuals or 10% of the sample, whichever came first. We used an ocular micrometer to measure the length of each individual to the nearest half increment (0.24 mm). Then we used length-mass regressions to convert length to dry mass, using equations for species with the most similar body shape when we were unable to identify the morphospecies to as low a taxonomic level as the length-mass regressions were organized. Specifically, we used Dumont et al. (1975) for *Moina* using the equation for *M. micrura*, and for Chydoridae using the equation for *Alonella exigua*; Benke et al. (1999) for insect larvae; McCauley (1984) for *Scapholeberis*, and for *Daphnia* using the equation for *D. longispina*; Culver et al. (1985) for Cyclopoida using the equation for nonovigerous female *Mesocyclops edax*; Baumgärtner and Rothhaupt (2003) for Hydracarina; Tellez et al. (2009) for Corixidae; and Caballero et al. (2004) for Collembola using the equation for *Folsomia candida.* We sorted the taxa into two non-mutually exclusive groups for analysis: zooplankton and benthic taxa.

### Chemical analyses

We weighed the dried algal mass samples, then split them into two subsamples, one of which we combusted using a muffle furnace for two hours at 550 °C. We weighed the resulting ash to calculate the ash-free dry weight. Separately, we added enough NaOH to the media samples to return the pH to 7. Then we digested all of these samples using a Hach Simplified TKN kit (s-TKN™) to convert all organic N and P to inorganic forms. Lastly we analyzed ammoniacal N, total Kjeldahl N (TKN), NO_x_ (summed nitrate and nitrite), total N (TN), and PO_4_ in all samples using a Hach spectrophotometer and Hach Ammonia Test Kit Model NI-8, the Hach Simplified TKN kit (s-TKN™), and the Hach Orthophosphate Test Kit Model PO-19A. *Data analysis*

To analyze the effects of the treatments on the dependent variables including zooplankton and herbivore mass, dry algal mass, algal biovolume, and nutrient concentrations, we fit and compared nested sets of models including and excluding treatment effects. First, we visually assessed the whether the residuals more closely followed a normal or a gamma distribution depending on the right skewedness. Based on this assessment we used linear models for the analyses of dry algal weight and of raw algal nutrient content variables, and we used gamma GLMs for the analyses of algal ash P content, algal biovolume, and invertebrate mass variables. We separately analyzed the effect of cage side and fish presence. Cage side is whether the cage was surrounding the paddlewheel or on the opposite side of the pond (Fig. 1), which determined how much of the pond the fish could access. The nested models were as follows: cage side + fish presence, fish presence, and the null model with intercept only. We did not include a random effect of tank row because our sample size of 18 did not provide enough power to include a random effect without overfitting the models. We assessed all model fits using plots of the residuals, then compared nested sets of GLMs and LMs with likelihood ratio tests and F-tests, respectively. Additionally, we created NMDS plots using the *vegan* package to assess the effect of the treatments on algal and invertebrate community composition, both including and excluding taxa only found in one sample (Oksanen et al. 2022). All analyses were performed in R version 4.2.2 (R Core Team 2022).

## Results

### Temperatures and fish

Unseasonably warm temperatures began on May 3 and continued to increase for the next two weeks, with daily highs above 30 °C and reaching 36 °C. Water temperatures rose as high as 34.1 °C on May 8, after which we began adding ice. Eight fish were replaced throughout the course of the experiment, most of which likely were succumbing to the high temperatures. As a result, following the IACUC’s recommendation we removed all fish and ended the experiment following the sampling on May 17.

### Invertebrates

The sampled zooplankton community was dominated by *Moina* (Online Resource 1 Table S2), while the benthic invertebrate community was dominated by small-bodied cladocerans of the family Chydoridae and larvae of the dipteran family Chironomidae (Online Resource 1 Table S3). Few if any *Streptocephalus* survived to the time of sampling and none were collected in samples. Zooplankton mass was 88.8% lower in the presence of fish; however, zooplankton mass was 93.1% higher when the cage surrounded the paddlewheel (Tables 1a, 2a). This meant that zooplankton mass was higher when fish had access to more area compared to tanks in which fish were confined to a cage (Fig. 2a). Benthic invertebrate mass was not affected by fish presence or the area fish could access (Table 1b; Fig. 2b). Overall invertebrate community composition did not appear to correlate with treatment, based on substantial overlapping in NMDS space (Online Resource 1 Figs. S1-S3). This result held when rare taxa were removed (Online Resource 1 Figs. S4-S6).

**Table 1.** Results of likelihood ratio tests and F-tests comparing nested GLMs and LMs, respectively, for a) zooplankton biomass, b) benthic invertebrate biomass, c) algal dry weight, d) algal biovolume, e) available ammoniacal N (concentration in the water), f) available NO_x_-N, g) available total Kjeldahl N, and h) available total N. The models including only the effect of fish presence were compared against the null models, and the full models were compared against the models with fish presence only. Presented are the model class (GLM = gamma GLM, LM = linear model), degrees of freedom (difference in the number of parameters, Df), deviance for GLMs (inverse goodness of fit), F value for LMs, and *p* value for the comparison (bolded if < 0.05).

| Model | Class | Df | Deviance | F | *p* |
| --- | --- | --- | --- | --- | --- |
| **a) zooplankton** |  |  |  |  |  |
| fish presence | GLM | 1 | 20.15 |  | **<0.001** |
| fish presence + caged paddlewheel | GLM | 1 | 10.13 |  | **0.013** |
| **b) benthic invertebrates** |  |  |  |  |  |
| fish presence | GLM | 1 | 0.4095 |  | 0.339 |
| fish presence + caged paddlewheel | GLM | 1 | 0.3411 |  | 0.383 |
| **c) algal dry weight** |  |  |  |  |  |
| fish presence | LM | 1 |  | 6.589 | **0.021** |
| fish presence + caged paddlewheel | LM | 1 |  | 0.481 | 0.499 |
| **d) algal biovolume** |  |  |  |  |  |
| fish presence | GLM | 1 | 1.376 |  | **0.029** |
| fish presence + caged paddlewheel | GLM | 1 | 0.3107 |  | 0.299 |
| **e) ammoniacal N** |  |  |  |  |  |
| fish presence | LM | 1 |  | 0.0556 | 0.817 |
| fish presence + caged paddlewheel | LM | 1 |  | 0.4665 | 0.505 |
| **f) NO<sub>x</sub>-N** |  |  |  |  |  |
| fish presence | LM | 1 |  | 2.115 | 0.167 |
| fish presence + caged paddlewheel | LM | 1 |  | 0.1696 | 0.686 |
| **g) TKN** |  |  |  |  |  |
| fish presence | LM | 1 |  | 0.0018 | 0.966 |
| fish presence + caged paddlewheel | LM | 1 |  | 4.353 | 0.054 |
| **h) Total N** |  |  |  |  |  |
| fish presence | LM | 1 |  | 0.0198 | 0.890 |
| fish presence + caged paddlewheel | LM | 1 |  | 4.140 | 0.060 |

**Table 2.**
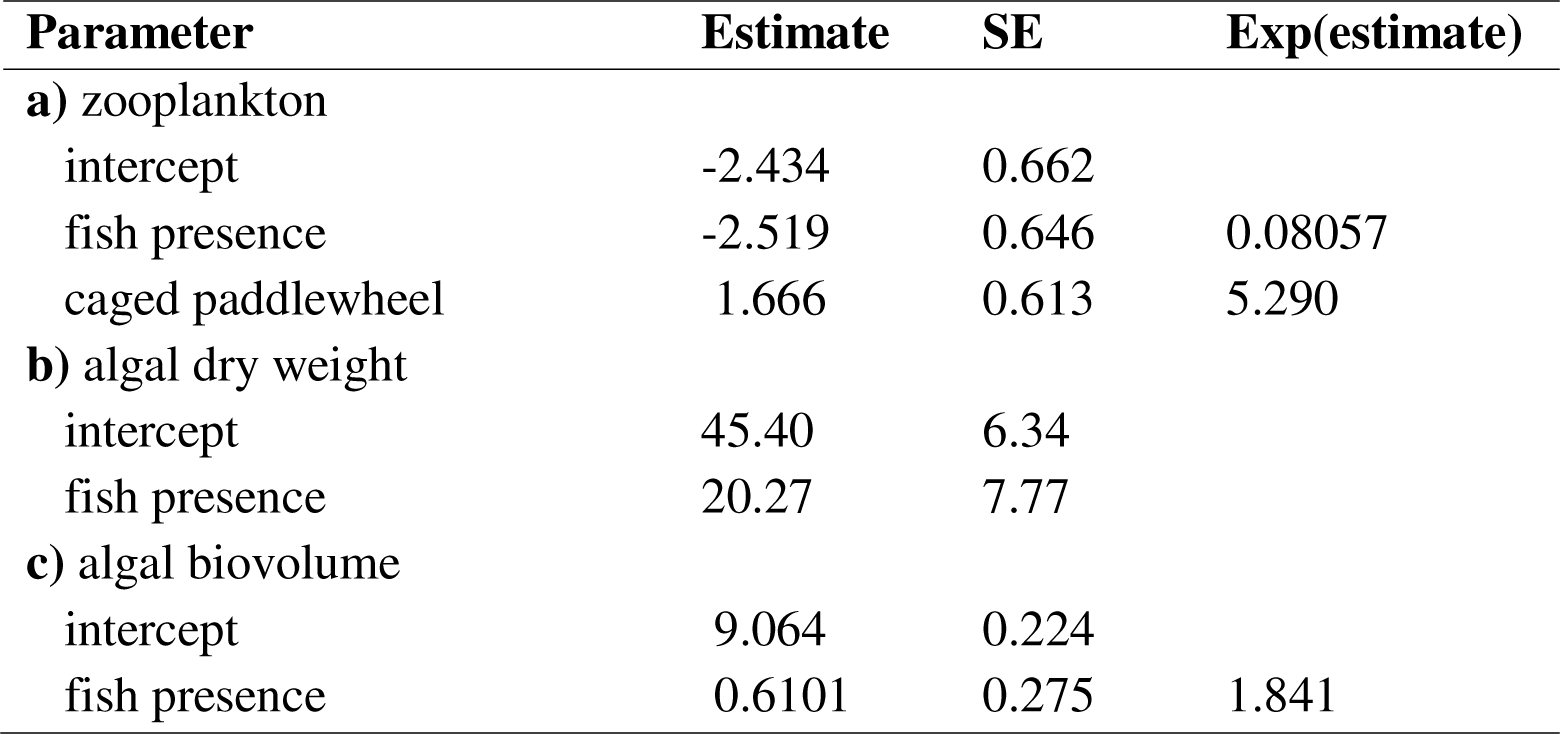
Coefficient estimates for the best models as determined by model comparisons (Table 1), for cases where the null model was not supported. Estimates are displayed along with their standard errors (SE) and, for GLMs, their natural exponential functions [‘‘Exp(estimate)’’], which can be interpreted as multiplicative effects (e.g., fish reduced zooplankton biomass to 0.08057× the biomass without fish).

| Parameter | Estimate | SE | Exp(estimate) |
| --- | --- | --- | --- |
| **a) zooplankton** |  |  |  |
| intercept | -2.434 | 0.662 |  |
| fish presence | -2.519 | 0.646 | 0.08057 |
| caged paddlewheel | 1.666 | 0.613 | 5.290 |
| **b) algal dry weight** |  |  |  |
| intercept | 45.40 | 6.34 |  |
| fish presence | 20.27 | 7.77 |  |
| **c) algal biovolume** |  |  |  |
| intercept | 9.064 | 0.224 |  |
| fish presence | 0.6101 | 0.275 | 1.841 |

**Figure 2.**
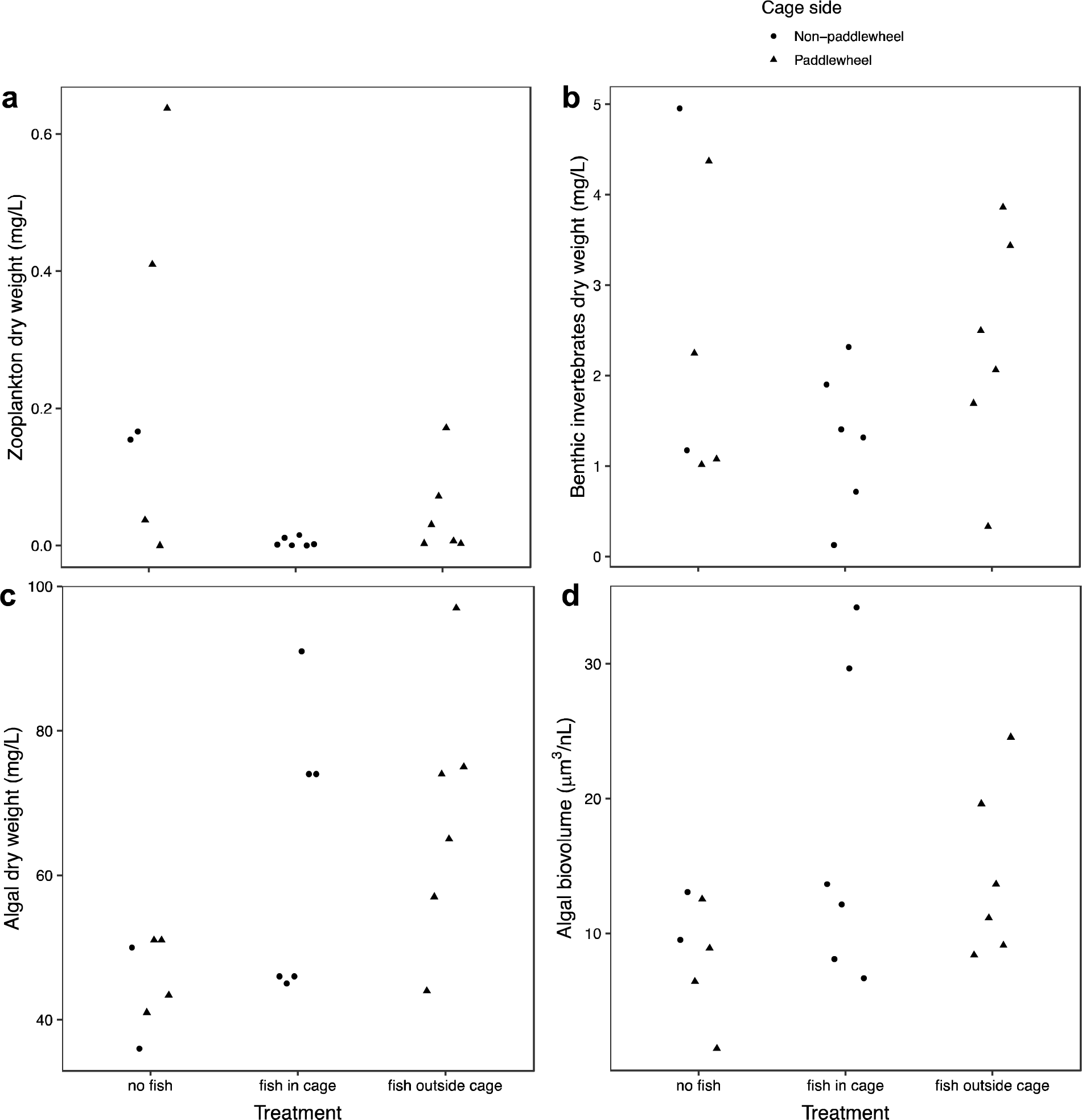
Invertebrate and algal mass metrics by fish access treatment and cage placement. **a**: Zooplankton dry weight; **b**: benthic invertebrate dry weight; **c**: algal dry weight; and **d**: algal biovolume

### Algae

By the time of sampling the algal community became dominated by *Actinastrum*, *Planktothrix*, *Micractinium*, and *Desmodesmus quadricauda*. Other less common morphospecies were mostly chlorophytes (Online Resource 1 Tables S1, S4). Sunfish enhanced dry algal weight by 44.7% and algal biovolume by 84.1%, but cage placement did not significantly affect either metric (Tables 1c-d, 2b-c; Fig. 2c-d). Algal community composition did not appear to correlate with treatment, whether including or excluding rare taxa, based on substantial overlapping in NMDS space (Online Resource 1 Figs. S7-S12).

### Nutrients

Concentrations of ammoniacal N, NO_x_, TKN, and TN in the water were not significantly affected by treatment (Table 1e-h; Fig. 3). Similarly, none of the algal nutrient content metrics were significantly affected by fish or cage placement (Online Resource 1 Table S5, Fig. S13).

**Figure 3.**
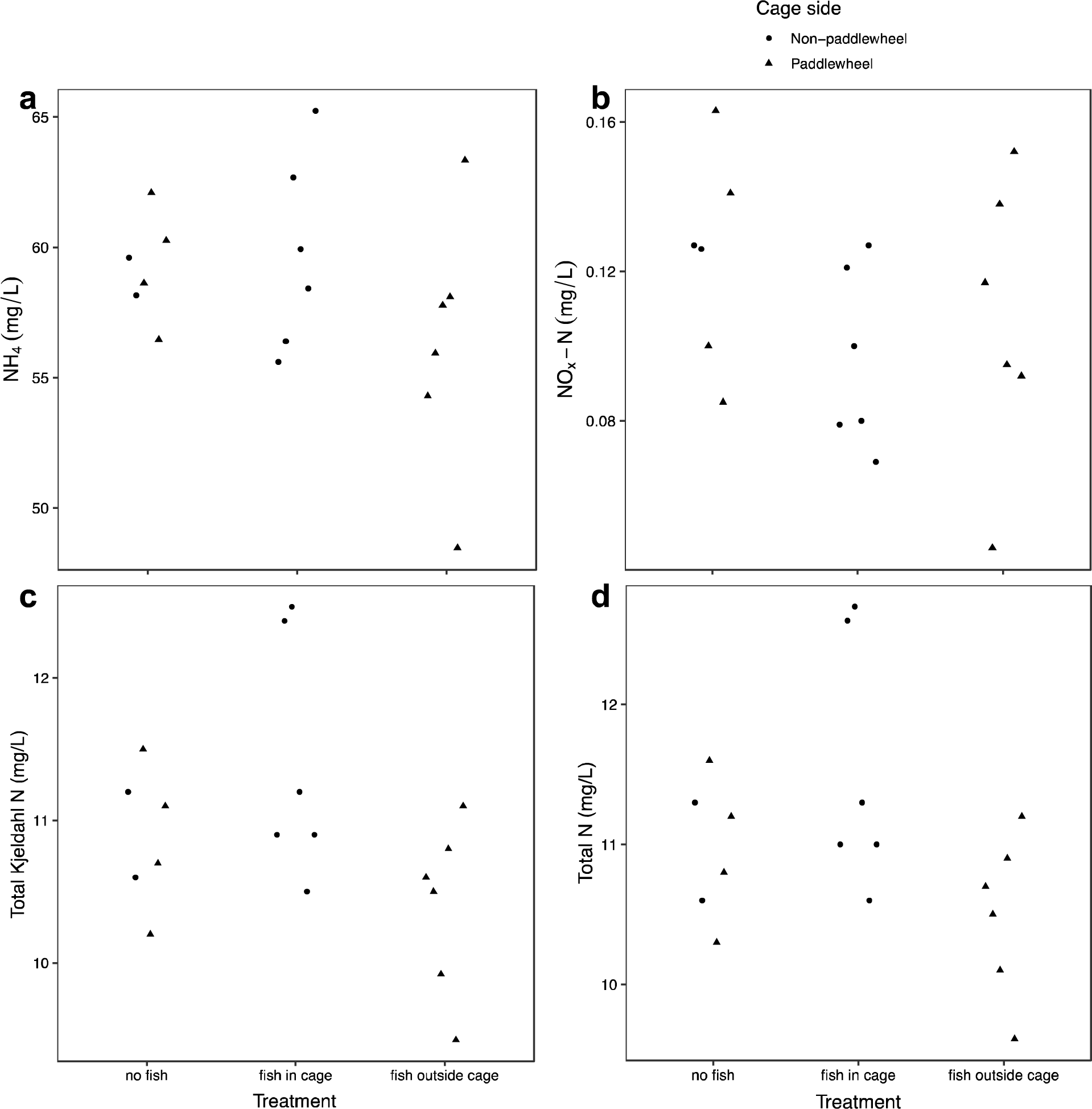
Dissolved nutrient metrics by fish access treatment and cage placement. **a**: ammonium, **b**: NO_x_-N, **c**: total Kjeldahl N (TKN), **d**: total N (TN)

## Discussion

We tested potential methods of biological control in WAC with the goal of improving the efficiency of this environmentally-promising practice. After only 19 days, bluegill sunfish drastically reduced zooplankton biomass, especially when confined to a cage. This reduction of zooplankton by fish led to a 45% and 84% increase in algal dry weight and biovolume, respectively, regardless of the cage placement. However, none of the nutrient metrics we measured, whether dissolved nutrients or algal nutrient content, were affected by our treatments by the end of the experiment. It is possible that given more time, though, the enhanced algal biomass in ponds with fish may have increased nutrient uptake rates and decreased available nutrients in the medium.

Overall, despite higher diversity and therefore more complex food web than most algal cultivation ponds which are typically dominated by a single species of algae, our results indicate a classic trophic cascade in which the sunfish suppressed herbivorous zooplankton enough to allow microalgae to reproduce faster than they could be consumed. However, one of the more interesting and surprising results involved the cage placement and invertebrate biomass. When the cage was surrounding the paddlewheel, which meant the fish were outside the cage when present, zooplankton biomass was higher than when the cage was opposite the paddlewheel. This result fit our prediction that when fish were outside the cage they would incorporate a higher proportion of fast and benthic prey into their diet, weakening their top-down effect on zooplankton. However, there is no evidence that benthic invertebrates were reduced in this or any treatment. Therefore, the association between cage placement and zooplankton might have resulted from another factor such as the cage providing some protection to the zooplankton by reducing damage from the paddlewheel, rather than an aspect of the food web. Future experiments will be needed to confirm the generality of these results.

Our results agree with the bulk of freshwater trophic cascade studies which show that predators often have strong effects on the biomass of lower trophic levels – the sign of which alternates for each successive lower trophic level – but that the effects attenuate down the food web (Brett and Goldman 1996; Borer et al. 2005). It makes intuitive sense that the more indirect an effect, the weaker and slower the effect will tend to be. For a biological control method to be maximally effective in a raceway WAC system, therefore, many algal generations may be needed for the trophic cascade to affect the available nutrients and allow enhanced nutrient uptake rates.

Using mosquitofish instead of sunfish, Sturm et al. (2012) found the same qualitative effects of fish on algal biomass and nutrient uptake in wastewater treatment ponds as we did. The duration of their experiment was >80 days, more than 4× the length of our fish deployment. After 25 days, they still did not see an effect of the fish on algal biomass, but by the end of the experiment after >80 days they saw an even stronger effect of fish on algal biomass than in our experiment: a 3-fold increase. However, even this large boost of algal biomass did not result in an increase in nutrient removal from the water.

We believe this study is useful as a starting point for biological control research in raceway WAC, but it is prudent to note a few limitations to our experiment. First, we were only able to leave the fish in the tanks for 19 days before high temperatures necessitated their removal. This relatively short duration leaves open the question of whether a longer incubation period would have allowed the trophic cascade to reach the dissolved nutrients and improve the nutrient removal potential of the raceway ponds. Several fish also had to be replaced during those 19 days, likely due to heat stress. It is likely that as these fish became stressed, they ceased feeding, and as there was only a single fish per tank, this may have weakened the top-down effect of the fish on the food web. The small size and relatively slow flow rate of the raceway ponds could have also led to different dynamics than would be observed in large WAC systems. For example, it is conceivable that a faster flow rate could make it more difficult for fish to prey on zooplankton, especially since fish cannot easily rely on visual cues in the turbid water of an algal culture. Lastly, this method is only appropriate for continuous cultivation in less toxic wastewaters such as agricultural runoff or pre-treated municipal wastewaters. Batch cultivation in which algae is grown to a target density, harvested in full, after which the pond is drained and re-inoculated, would not be easily compatible with a self-sustaining predator population, although it may be possible to temporarily move the fish or other predators to another pond before returning them following re-inoculation. Highly toxic wastewaters would not be appropriate for even the hardiest of fish, although some predaceous protozoans such as amoeba may be candidates for biological control agents in such systems. However, such toxic wastewaters are inhospitable to most zooplankton as well, and even most algae, meaning there is less need for zooplankton control in these systems.

Hardy zooplanktivorous fish like bluegill appear to represent a promising strategy for enhancing algal production in relatively benign wastewater such as agricultural runoff or tertiary municipal wastewater. As raceway ponds are shallow and operated in full sunlight, fish will be most likely to thrive in these ponds in cooler climates. In subtropical and tropical climates, other more heat-tolerant predaceous taxa such as protozoans like *Amoeba* or insects like pleids would be good candidates for future experimentation. Implementation of biological control in algal cultivation would represent relatively little economic or environmental cost, but could significantly improve the rate at which microalgae convert polluting nutrients to harvestable biomass. This algal biomass can be used as a biofuel feedstock, organic fertilizer, or other uses and generate revenue to maintain the cultivation operation. Therefore, coupled WAC systems could simultaneously combat several pressing environmental challenges intertwined in the water-food-energy security nexus upon which the wellbeing of humanity and the planet hang in the balance.

## Supporting information

Online Resource 1

## Funding

This research was supported by the Department of Integrative Biology at the University of Texas at Austin.

## Acknowledgements

A special thank you to Hans Hofmann for his guidance in navigating the IACUC protocol approval and revision process. Thank you to Chris Dutton for his generosity in sharing his knowledge and 3D printer files for constructing the paddlewheels; Nhan Tran, Nikki Hy, and Ishita Kansal who helped set up the experiment and complete laboratory work; Caroline Farrior and Shalene Jha who provided advice and feedback on the manuscript; Isaac Miller-Crews who lent us equipment, and Sisimac Duchicela who assisted in the field; and Stephen Peña, Rakib Rashel, and Matt Ashworth who provided assistance in the laboratory. This research was made possible by Brackenridge Field Laboratory including help from Jason Lawson.

## Notes

### Competing Interest Statement

The authors have declared no competing interest.

