## Supplementary material for "Sunfish as zooplankton control agents to improve yields of wastewater-cultivated algae": Online Resource 1

**Table S1.** Cell shapes and mean cell biovolumes of algal taxa/morphospecies. Taxa are listed in descending order of total biovolume across all raceway ponds.

*Identified using molecular methods (see Table S4).

| **Taxon** | **Shape** | **Mean cell biovolume (μm^3^)** | **Notes** |
| --- | --- | --- | --- |
| *Actinastrum* | ½ cone and ½ cylinder | 30.1 |  |
| *Planktothrix* | cylinder | 3.08 |  |
| *Micractinium** | sphere | 19.7 |  |
| *Desmodesmus quadricauda* | prolate spheroid | 8.43 |  |
| Sphaeropleales* | ovoid | 53.3 | solitary; possibly *Scenedesmus obliquus* |
| *Scenedesmus* sp. | sphere | 8.89 | colonial |
| *Nitzschia* | pennate | 207 |  |
| green picoplankton | sphere | 1.10 |  |
| *Selenastrum* | lunate | 35.5 |  |
| Oscillatoriales | cylinder | 2.81 | possibly *Arthrospira* |

**Table S2.** Zooplankton taxa in descending order of mean biomass across all raceway ponds.

| **Taxon** | **Classification** | **Mean mass (μg/L)** |
| --- | --- | --- |
| *Moina* | Cladocera | 93.9 |
| Elmidae larva | Insecta | 0.791 |
| *Scapholeberis* | Cladocera | 0.398 |
| *Daphnia* | Cladocera | 0.270 |

**Table S3.** Benthic invertebrate taxa in descending order of mean biomass across all raceway ponds.

| **Taxon** | **Classification** | **Mean mass (μg/L)** |
| --- | --- | --- |
| Chydoridae | Cladocera | 941 |
| Chironomidae larva | Insecta | 564 |
| Ephemeroptera nymph | Insecta | 340 |
| Corixidae | Insecta | 80.2 |
| Diptera larva | Insecta | 49.9 |
| Chironomidae pupa | Insecta | 19.4 |
| Collembola | Hexapoda | 1.23 |
| Hydracarina | Arachnida | 0.923 |

**Table S4.** DNA barcoding matches and primers used.

| **Taxon** | **BLAST** | **% Identity** | **Primer** |
| --- | --- | --- | --- |
| *Micractinium* | *Micractinium* sp. | 89.97 | ITS2 forward |
| *Micractinium* | *Micractinium* sp. | 99.78 | ITS2 reverse |
| Sphaeropleales | *Tetraedron* sp. | 98.41 | 23S forward |
| Sphaeropleales | *Tetraedron* sp. | 98.47 | 23S reverse |

**Table S5.** Results of F-tests and likelihood ratio tests comparing nested LMs and GLMs, respectively, for algal nutrient content: a) total P of raw algae, b) total P of algal ash, c) NO_x_-N of raw algae, d) total Kjeldahl N of raw algae, and e) total N of raw algae. The models including only the effect of fish presence were compared against the null models, and the full models were compared against the models with fish presence only. Presented are the model class (GLM = gamma GLM, LM = linear model), degrees of freedom (difference in the number of parameters, Df), deviance for GLMs (inverse goodness of fit), F value for LMs, and *p* value for the comparison.

| **Model** | **Class** | **Df** | **Deviance** | **F** | ***p*** |
| --- | --- | --- | --- | --- | --- |
| **a)** TP raw algae |  |  |  |  |  |
| fish presence | LM | 1 |  | 0.5338 | 0.476 |
| fish presence + caged paddlewheel | LM | 1 |  | 0.0161 | 0.901 |
| **b)** TP algal ash |  |  |  |  |  |
| fish presence | GLM | 1 | 0.5207 |  | 0.306 |
| fish presence + caged paddlewheel | GLM | 1 | 0.3479 |  | 0.403 |
| **c)** NO_x_-N raw algae |  |  |  |  |  |
| fish presence | LM | 1 |  | 0.0285 | 0.868 |
| fish presence + caged paddlewheel | LM | 1 |  | 2.319 | 0.149 |
| **d)** TKN raw algae |  |  |  |  |  |
| fish presence | LM | 1 |  | 0.0248 | 0.877 |
| fish presence + caged paddlewheel | LM | 1 |  | 0.2362 | 0.634 |
| **e)** TN raw algae |  |  |  |  |  |
| fish presence | LM | 1 |  | 0.0254 | 0.876 |
| fish presence + caged paddlewheel | LM | 1 |  | 0.2201 | 0.646 |

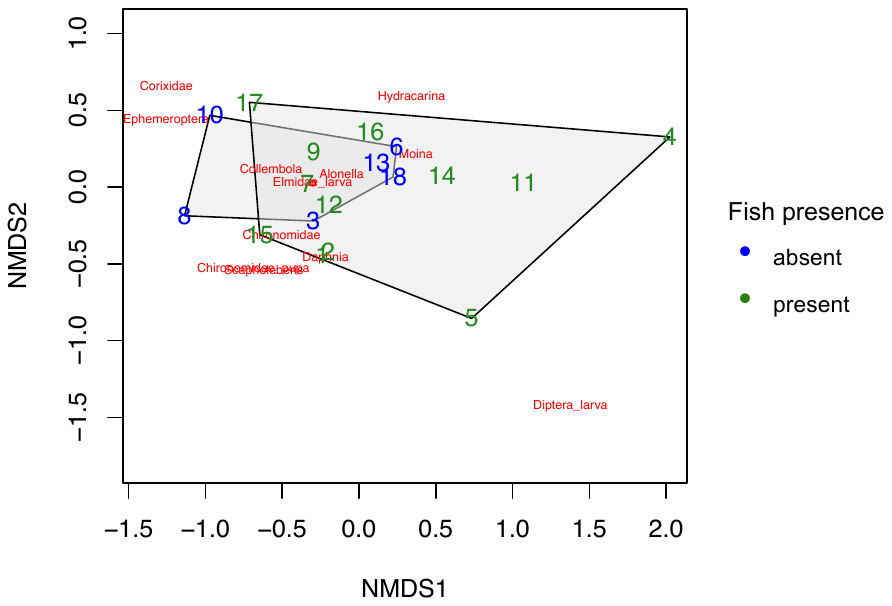

**Figure S1.** NMDS plot of invertebrate community composition by fish presence. Numbers represent tanks.

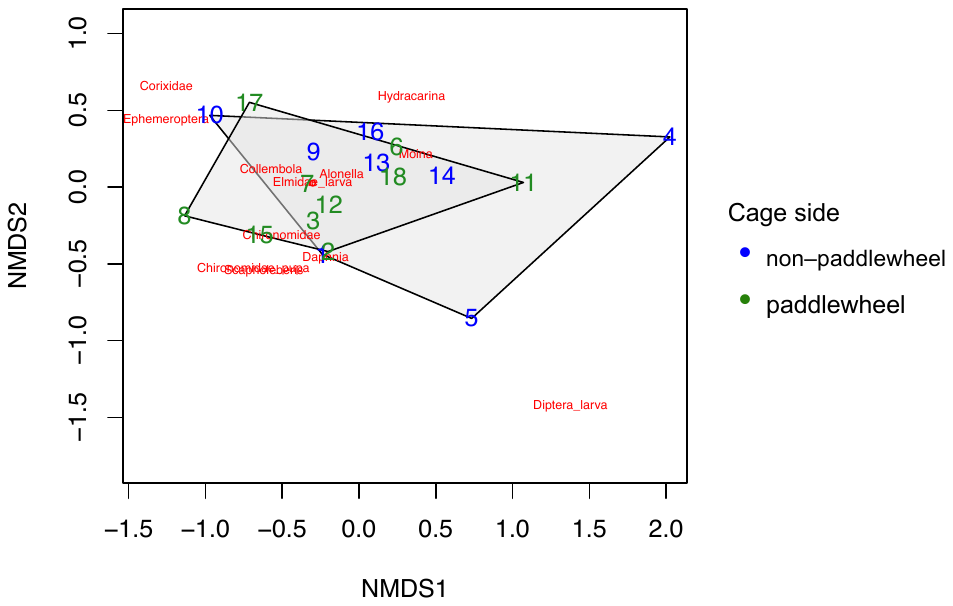

**Figure S2.** NMDS plot of invertebrate community composition by cage placement (on the opposite side from the paddlewheel or surrounding the paddlewheel). Numbers represent tanks.

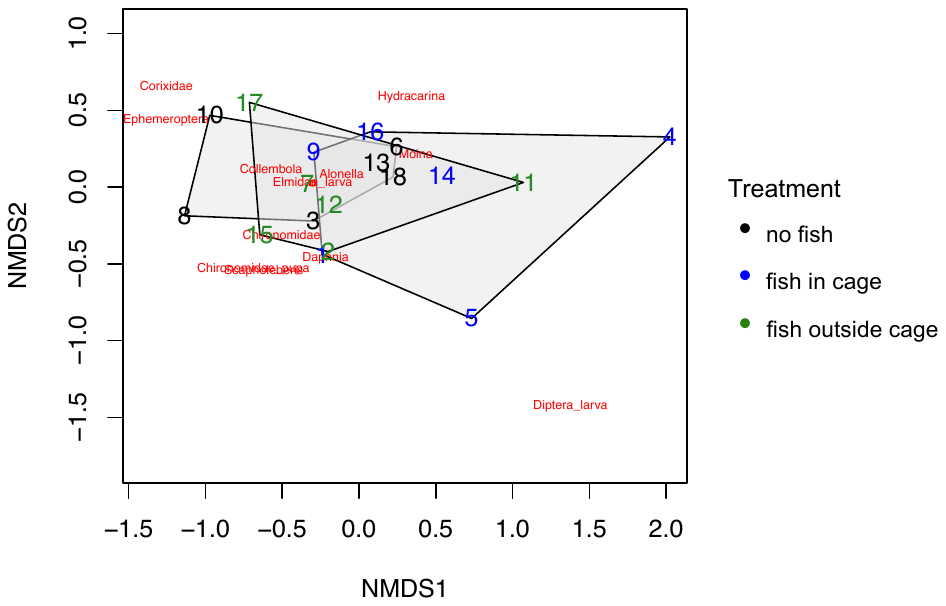

**Figure S3.** NMDS plot of invertebrate community composition by treatment (no fish, fish inside cage, fish outside cage). Numbers represent tanks.

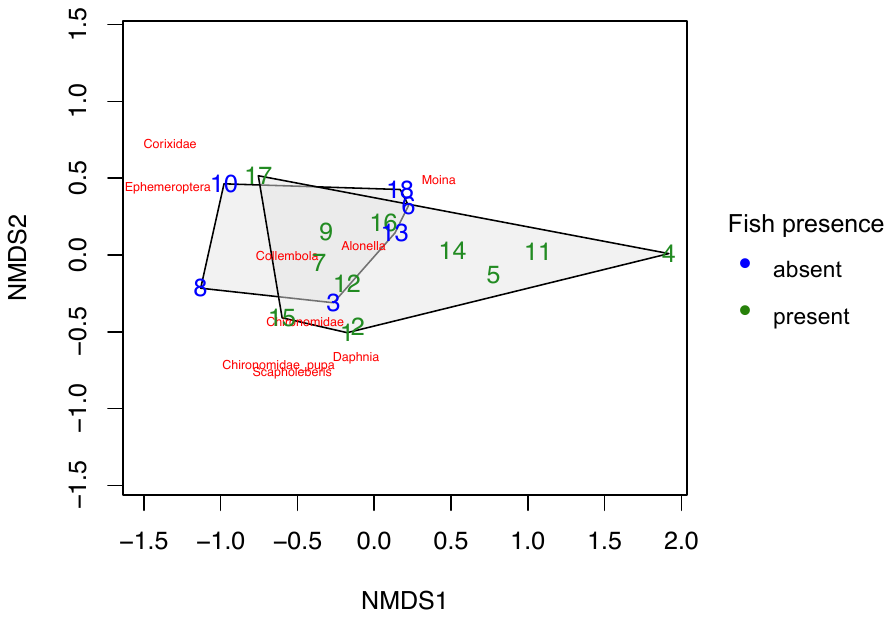

**Figure S4.** NMDS plot of invertebrate community composition by fish presence, excluding taxa found in only one sample. Numbers represent tanks.

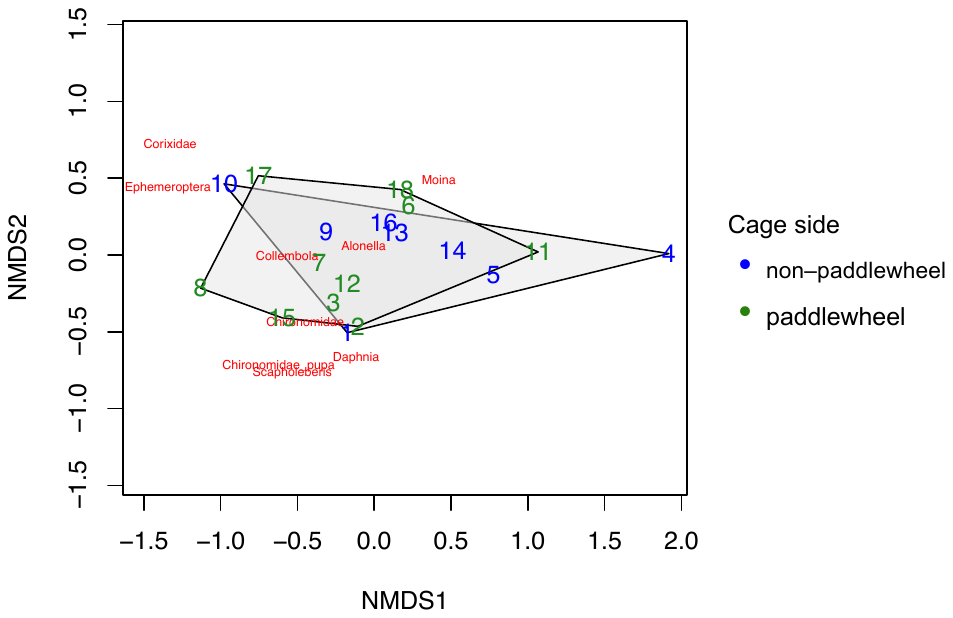

**Figure S5.** NMDS plot of invertebrate community composition by cage placement (on the opposite side from the paddlewheel or surrounding the paddlewheel), excluding taxa found in only one sample. Numbers represent tanks.

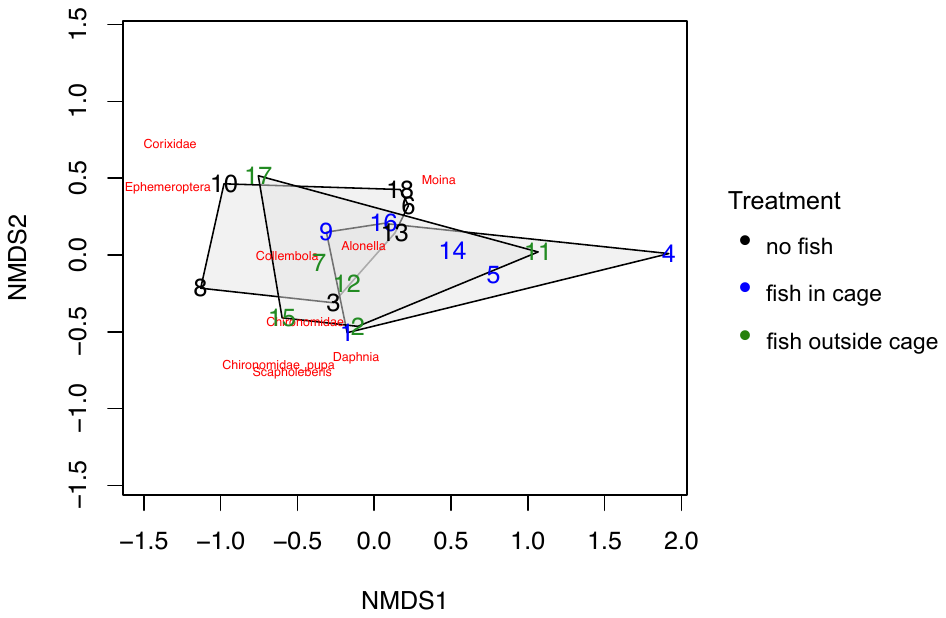

**Figure S6.** NMDS plot of invertebrate community composition by treatment (no fish, fish inside cage, fish outside cage), excluding taxa found in only one sample. Numbers represent tanks.

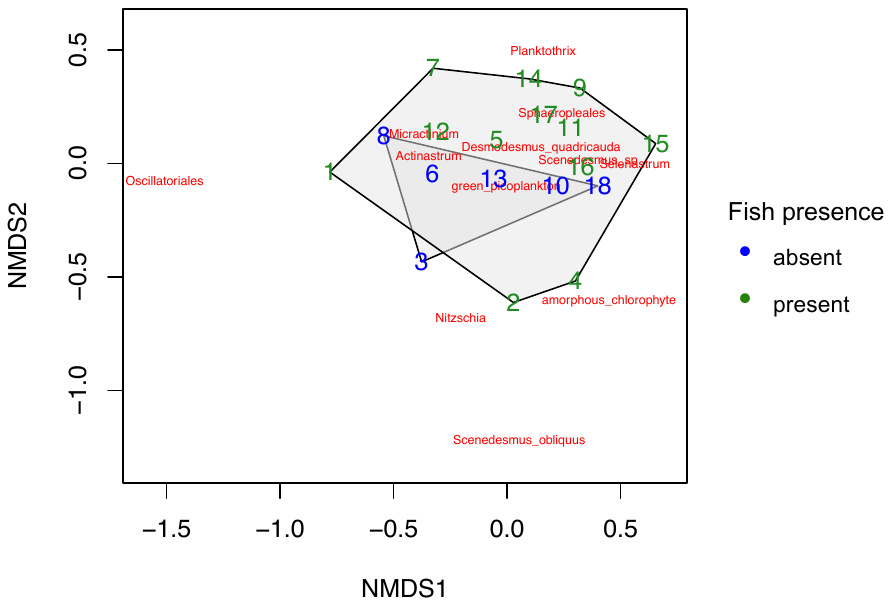

**Figure S7.** NMDS plot of algal community composition by fish presence. Numbers represent tanks.

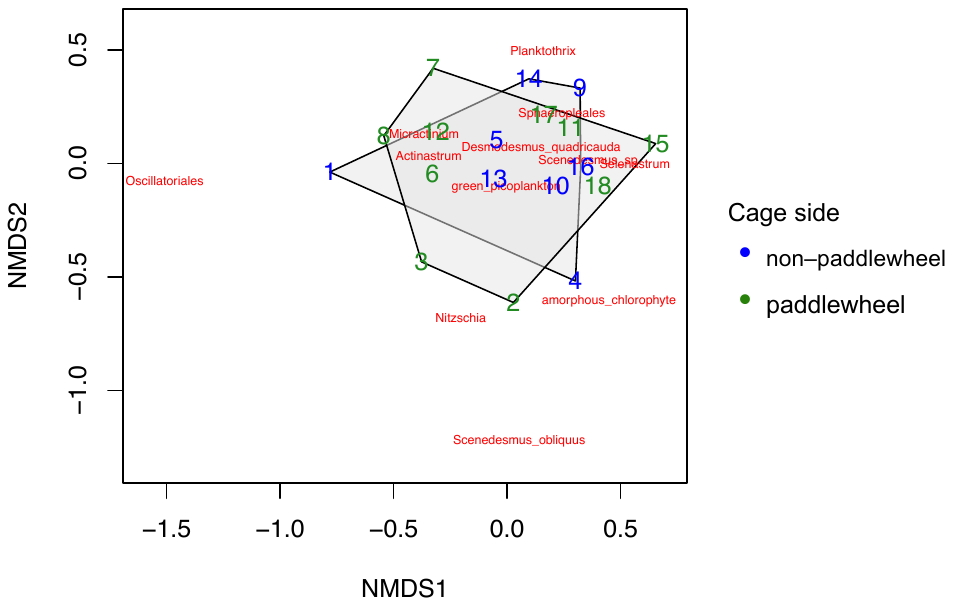

**Figure S8.** NMDS plot of algal community composition by cage placement (on the opposite side from the paddlewheel or surrounding the paddlewheel). Numbers represent tanks.

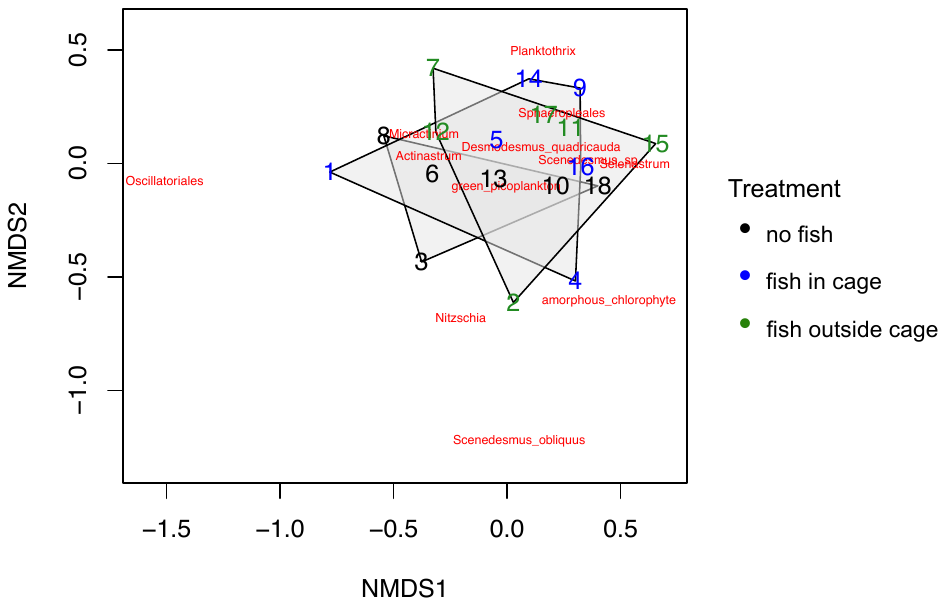

**Figure S9.** NMDS plot of algal community composition by treatment (no fish, fish inside cage, fish outside cage). Numbers represent tanks.

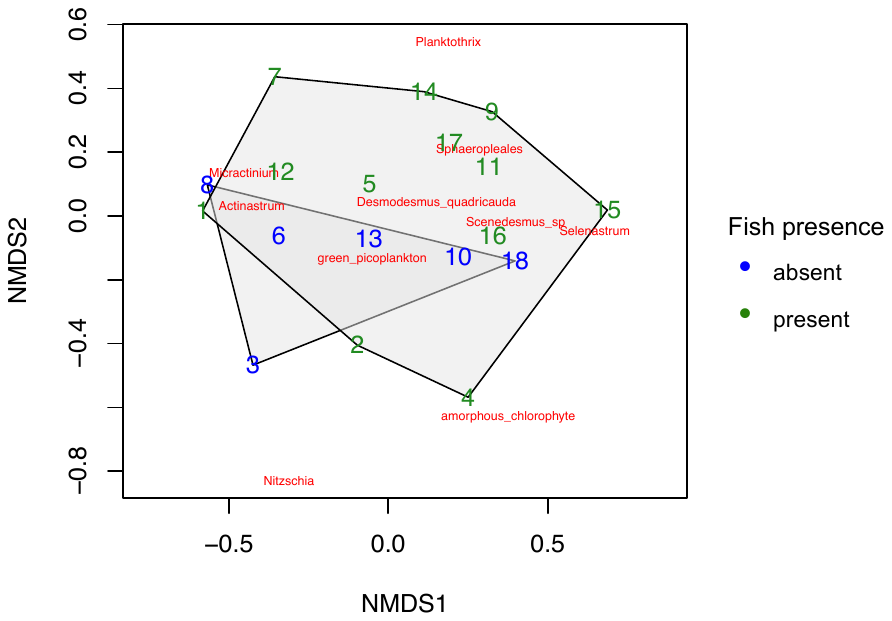

**Figure S10.** NMDS plot of algal community composition by fish presence, excluding taxa found in only one sample. Numbers represent tanks.

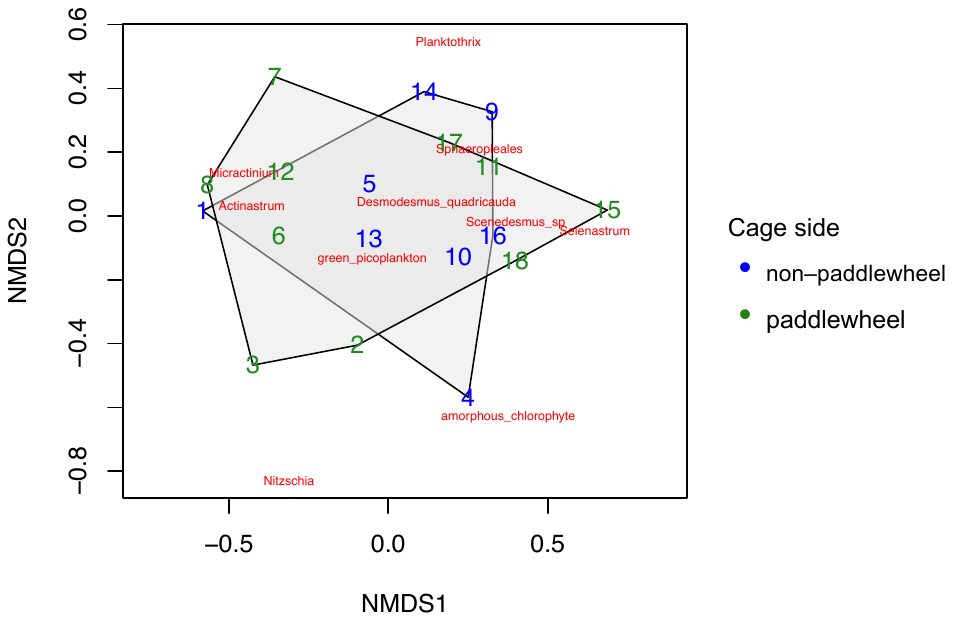

**Figure S11.** NMDS plot of algal community composition by cage placement (on the opposite side from the paddlewheel or surrounding the paddlewheel), excluding taxa found in only one sample. Numbers represent tanks.

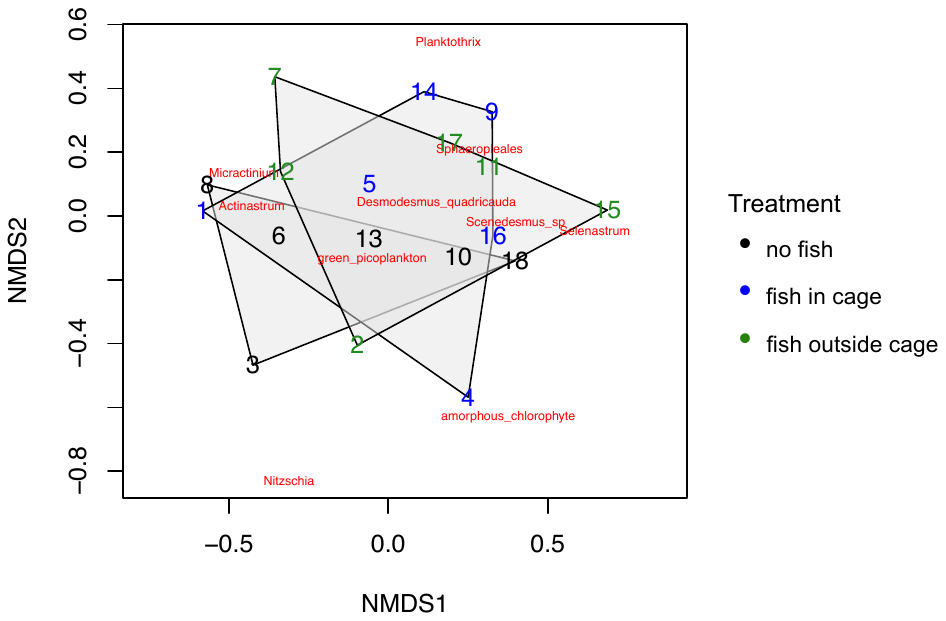

**Figure S12.** NMDS plot of algal community composition by treatment (no fish, fish inside cage, fish outside cage), excluding taxa found in only one sample. Numbers represent tanks.

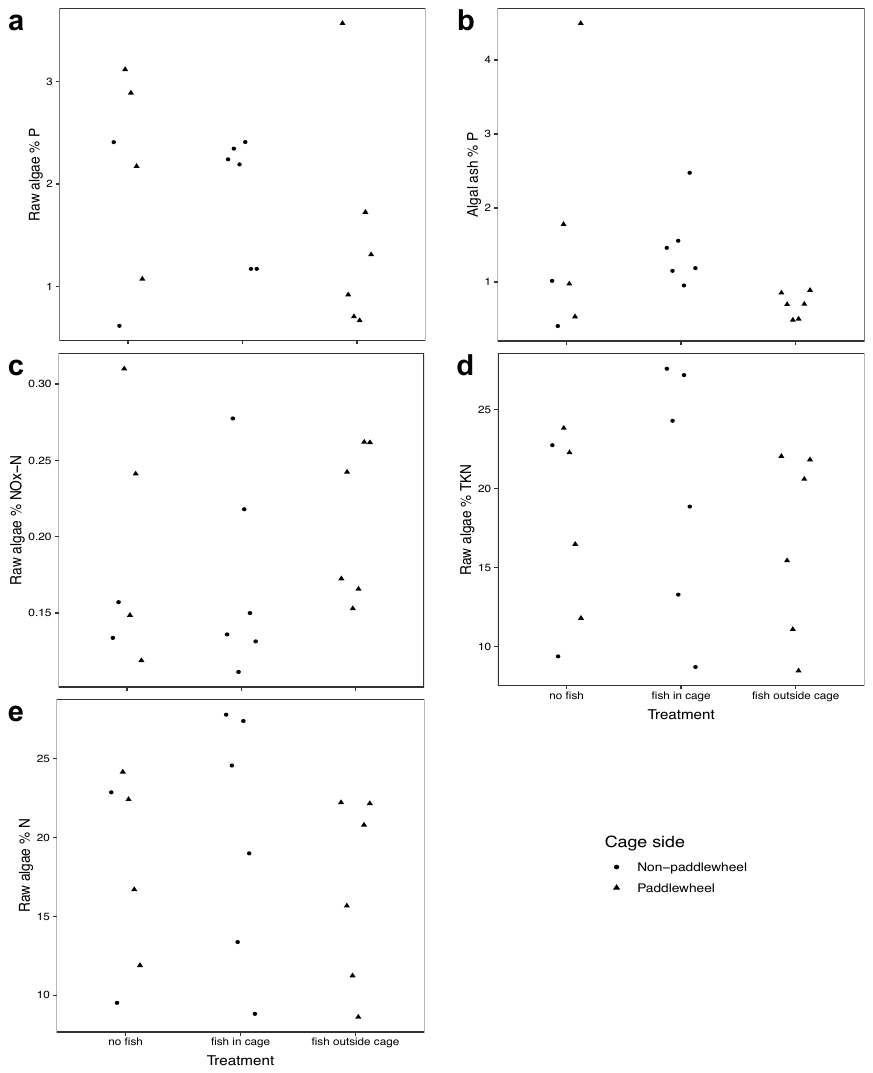

**Figure S13.** Algal nutrient content metrics by fish and cage placement. **a**: % P of raw algae, **b**: % P of algal ash, **c**: % NO_x_-N of raw algae, **d**: % total Kjeldahl N of raw algae, **e**: % N of raw algae.
